# Engineering *Saccharomyces cerevisiae* for the production of punicic acid-rich yeast biomass

**DOI:** 10.1101/2024.06.20.599979

**Authors:** Juli Wang, Guanqun Chen

## Abstract

Punicic acid (PuA), an unusual conjugated linolenic acid found in pomegranate, offers diverse health benefits and potential applications in the food industry. The limited production of PuA from its natural plant sources has led to a growing interest in producing it through microbial fermentation. In this study, the “generally recognized as safe” yeast *Saccharomyces cerevisiae* was engineered to produce PuA. Candidate genes related to PuA synthesis were directly shuffled within the yeast genome. Subsequent screening and optimization resulted in a strain that synthesized 26.7% of its total fatty acids as PuA. Further analyses revealed that the strain’s storage lipids contained over 22% PuA. By incorporating the health-promoting lipid component into the nutritional profile of *S. cerevisiae*, the engineered strain could potentially serve as a sustainable source of yeast biomass with enhanced nutritional value.

## 1. Introduction

Pomegranate (*Punica granatum*) derived punicic acid (PuA; 18: 3Δ^9*cis*,11*trans*,13*cis*^) is a typical conjugated linolenic acid and a representative plant-derived unusual fatty acid. ^1^ The three conjugated double bonds in PuA offer a wide range of bioactivities. ^2–4^ Therefore, both PuA and pomegranate seed oil have been studied extensively for their potential health benefits and could be used as functional ingredients in food and nutraceuticals. ^3–10^ Due to these emerging applications, the potential market of PuA is expected to experience significant growth in the coming years. ^1,2^ As a result, the market price of pomegranate seed oil, depending on its purity, has been reported to be relatively high. ^1,2,11^ However, growing pomegranate for large-scale PuA production has significant challenges, such as a low seed-to-fruit ratio and low oil yield compared to oilseed crops. ^12,13^ In this regard, metabolic-engineered microorganisms emerge as a promising alternative.

For food and feed use, metabolic engineering efforts have led to significant advancements in producing yeast biomass and single-cell oil enriched with healthy fatty acids. One notable example is the production of eicosapentaenoic acid (EPA) in the oleaginous yeast *Yarrowia lipolytica*. ^14–16^ Through the integration of thirty copies of genes involved in the fatty acid elongation and desaturation pathways, DuPont Applied BioSciences has engineered the yeast cell to produce high levels of EPA. ^14,15^ The resulting EPA-rich oil from yeast biomass has been approved by the US Food and Drug Administration (FDA) for food use and is marketed as New Harvest EPA oil for nutritional supplements. ^16,17^ Additionally, the EPA-enriched yeast biomass has found applications in aquaculture, such as feeding salmon under the brand name Verlasso to enhance the nutritional profile of the farmed fish. ^16,18^

Additionally, producing polyunsaturated and conjugated fatty acids embedded within yeast biomass may also contribute to the maintenance of product stability. While these fatty acids offer numerous health benefits, they are inherently more susceptible to oxidation because of their high degree of unsaturation and the conjugated configuration of double bonds. The oxidation can result in quality deterioration and the development of off-flavors in the oil product. ^19,20^ In this context, embedding health-promoting fatty acids in yeast biomass may prove advantageous since it has been shown that polyunsaturated or conjugated fatty acid-rich oils encapsulated within yeast biomass remain effectively shielded from oxidation for more than seven weeks, in contrast to extracted oils that began oxidizing almost immediately. ^21^

In terms of yeast species, *Saccharomyces cerevisiae*, commonly known as baker’s yeast, is the most widely used yeast in food and feed industries due to its “Generally Recognized as Safe” (GRAS) status and long history in food products. ^22,23^ Rich in protein and high in nutritional value, *S. cerevisiae* is applied extensively for producing single-cell proteins from cost-effective feedstocks. ^24,25^ In addition, the global demand for protein has driven many studies to explore health-promoting additives for use in animal feeds. In this regard, *S. cerevisiae* biomass has also emerged as a promising choice. ^26,27^ Supplementing animal feed with *S. cerevisiae* has been shown to improve animal health and performance, enhance feed efficiency and digestibility, and potentially reduce the negative effects of pathogenic bacteria. ^28–32^

Therefore, this study aims to develop biologically safe *S. cerevisiae* biomass containing PuA, the health-promoting conjugated linolenic acid derived from pomegranate. By targeting the locations of baker’s yeast Ty retrotransposon, genes that potentially contribute to PuA synthesis were shuffled on the yeast genome and led to a recombinant yeast library with various contents of PuA. Subsequent screening of the library as well as preliminary optimization led to a recombinant yeast strain with 26.7% of total fatty acids as PuA. In shake flask cultivation, the PuA titer reached 425 mg/L. To the best of our knowledge, the PuA content and titer are the highest achieved in engineered *S. cerevisiae*. The resulting yeast strain offers a potential sustainable source for producing nutritional yeast biomass, enhanced by the additional benefits of healthy conjugated linolenic acid.

## 2. Materials and methods

### 2.1. Strains, genes, and plasmids

All strains used in this study are listed in Table S1. In brief, *Escherichia coli* DH5α was used for routine plasmid construction and preparation. *S. cerevisiae* BY4741-*snf2*Δ obtained from the Euroscarf collection was used as the starting strain in this study. *S. cerevisiae* H1246 quadruple mutant lacking lipid biosynthetic ability was used to test the performance of pomegranate genes. Yeast transformations were performed using the traditional lithium acetate and PEG3350 method as described previously. ^33^ To obtain the template for cloning triacylglycerol (TAG) assembly and acyl-editing genes from *P. granatum*, total RNA was isolated from *P. granatum* tissues using the Spectrum Plant Total RNA Kit (Sigma-Aldrich, Oakville, Canada) and cDNA was synthesized using the SuperScript IV first-strand cDNA synthesis kit (Invitrogen, Carlsbad, USA). Putative sequences were obtained by performing an NCBI Basic Local Alignment Search Tool (BLAST) search against the online draft genome of *P. granatum*. ^34^

The target Ty retrotransposon region was analyzed based on the S288C reference genome from the *Saccharomyces* Genome Database (SGD) (http://www.yeastgenome.org/). Similarity and identity searches for target sequences in Ty retrotransposons were conducted using BLAST. Plasmids providing donor DNA were derived from pUC19. Four components, including upstream homology sequence, gene expression cassette, selection marker with truncated promoter region (*leu2*, *his3*, and *ura3* for round 1, 2, and 3 integrations, respectively), and downstream homology sequence were inserted. To release donor DNAs from the pUC19 backbone, either PCR or double digestion of SmaI sites flanking the donor DNA was conducted. For different genes, 500 fmol of each donor DNA were pooled together. To facilitate integration, CRISPR-Cas9 expression vectors (based on plasmid pCRCT, obtained from Addgene, plasmid #60621) targeting different locations on Ty retrotransposons were constructed and co-transformed into yeast cells with donor DNA. ^35^ Three 20bp sequences that are adjacent to the PAM sites that occur frequently in the respective target regions were selected as the spacer sequences for delivering crRNA (Table S2). These sequences were ordered as single-stranded, complementary oligos and annealed to obtain double-stranded DNAs. By adding homology arms, these sequences were inserted between two Eco31I restriction sites on the pCRCT plasmid.

### 2.2. Culture conditions and optimization

For plasmid construction and preparation, *E. coli* was cultured at 37°C in the Luria-Bertani (LB) medium with shaking at 225 rpm. To sustain the plasmids in *E. coli*, 100 mg/L ampicillin was applied. For testing PuA accumulation in shake flask cultivation, individual colonies of transformed yeast cells were first grown in 5 mL yeast nitrogen base (YNB) supplemented with an appropriate amino acid drop-out mix and 2% glucose for 24 h at 30°C with shaking. Seed cultures were then inoculated into 1 L shake flasks containing 60 mL synthetic complete medium (6.7 g/L YNB, 2 g/L synthetic complete supplement mixture of amino acids, and 6% glucose). The inoculum for PuA production experiments was grown for 72h at 20°C or 30°C, using an incubated shaker with a shaking speed of 250 rpm. For culture condition optimization, the Box– Behnken design was employed for response surface methodology. The factors considered for optimization were examined at three levels (6%, 10.5%, and 15% for glucose; 5.3, 6.5, and 7.6 for initial pH; 0.2, 0.6, and 1 for initial OD_600_). The two responses considered for analysis were PuA content (% of total fatty acids) and PuA titer (mg/L). Based on the results, second-order polynomial equations were fitted to predict the optimal points within experimental constraints (Table S3).

### 2.3. Preparation of plant oil hydrolysate and fatty acid feeding

The preparation of free fatty acids from pomegranate seed oil was conducted using chemical hydrolysis. ^36^ In brief, 50 g of pomegranate seed oil was mixed with 12 g potassium hydroxide, 117 mL pure ethanol, and 35 mL H_2_O in a shake flask flushed with nitrogen. The butylated hydroxytoluene (BHT) solution was added to the mixture to protect PuA. The reactor was then sealed and maintained at 50°C with constant shaking. After 1 h of incubation, removal of the unsaponifiable matter was performed three times by mixing 100 mL of distilled water and 100 mL of hexane. Free fatty acids were extracted with 100 mL hexane and dried over anhydrous sodium sulfate. The solvent was then removed under vacuum to obtain pomegranate oil hydrolysate. When fatty acid feeding was required, 0.03% v/v ethanol dissolved LA or pomegranate oil hydrolysate was supplemented to the culture medium along with 0.2% non-ionic surfactant NP-40 (TERGITOL™ solution) for even distribution of fatty acid in the aqueous medium.

### 2.4. Nile red staining of neutral lipids in yeast

Nile red fluorescence detection was conducted as described previously. ^37^ Briefly, 100 µl aliquots of the yeast cell suspension were transferred to a 96-well dark flat-bottom plate. Background fluorescence was measured using a Synergy H4 Hybrid multimode microplate reader (Biotek Instrument, Inc.) with emission and excitation filters set to 485 and 538 nm, respectively. A newly prepared methanolic Nile red solution (0.1 mg/mL) was added, and the second fluorescence intensity was measured. The Nile Red values were calculated based on the change in fluorescence over OD_600_ (ΔF/OD_600_).

### 2.5. Lipid extraction and separation of lipid class using thin-layer chromatography

Before lipid extraction, yeast biomass was harvested from liquid culture via centrifugation. The supernatant was then removed and 800 µL of a cold lipid extraction mixture comprising chloroform and isopropanol (2:1, v/v) was added, along with glass beads (0.5mm) and BHT at a final concentration of 0.01%. ^38,39^ Subsequently, cellular disruption was achieved through three cycles of bead beating (1-minute duration each) using a Biospec bead beater (Biospec, Bartlesville, OK), with 2-minute cooling on ice between each cycle. The extraction procedure was repeated twice for each sample. The collective organic phase, containing both polar lipid (PL) and TAG, was dried under nitrogen and resuspended in 200 µl chloroform. For lipid class analysis, the lipid extracted from each sample was separated using thin layer chromatography (TLC) with silica gel-coated plates (0.25 mm Silica gel, DCFertigplatten, Macherey-Nagel, Germany). The TLC plates were developed using a solvent mixture comprising hexane/diethyl ether/acetic acid (in a 70:30:1 ratio). ^40^ Lipid fractions on the TLC plate were visualized via 0.05% primulin staining under UV light. Bands corresponding to target lipid fractions were scraped, extracted, derivatized, and subsequently analyzed.

### 2.6. Positional analysis of triacylglycerol and polar lipids

The fatty acid distribution between *sn*-2 and *sn*-1/3 TAG was analyzed using previously described enzymatic reactions. ^40,41^ Briefly, after TLC separation, TAG was recovered from the silica gel, and dried under nitrogen. Subsequently, 1 mL Tris-HCl buffer (1 mM, pH 8.0), 100 μL 2.2% CaCl_2,_ and 250 μL 0.1% deoxycholate was added. Each mixture was vortexed for 2 minutes and sonicated for 60 seconds to emulsify the lipid. The mixture was pre-warmed in a water bath at 40°C for 30 s, and then 20 mg pancreatic lipase (pancreatic lipase type II, Sigma) was added to initiate hydrolysis. The mixture was further incubated for 3 min at 40°C, and the reaction was terminated by adding 500 μL of 6 M HCl. The resulting lipids, containing unreacted TAG, diacylglycerol (DAG), monoacylglycerol (MAG), and free fatty acids, were extracted twice with 3 mL of diethyl ether and then separated using TLC. The *sn*-2 MAG was subsequently analyzed.

The positional analysis of PL was performed by cleaving the fatty acids at the *sn*-1 position of PL using phospholipase A_1_. ^40,42^ In brief, phospholipase A_1_ was first mixed with water in a 1:4 (v/v) ratio. PL was recovered from the silica gel and dissolved in 2 mL of diethyl ether, and 1 mL of phospholipase A_1_ (Sigma) solution was added to initiate hydrolysis. The mixture was then vortexed at maximum speed for 5 min and the reaction was terminated by evaporation of diethyl ether under nitrogen. The hydrolyzed lipids were extracted and separated by TLC, and cleaved fatty acids were subsequently analyzed.

### 2.7. Lipid transmethylation and analysis

To prepare fatty acid methyl esters (FAMEs), transmethylation was carried out via a base-catalyzed method using 1 mL of 5% sodium methoxide dissolved in methanol. ^38,39^ After incubation at 30°C for 1 h, the reaction was stopped by adding 1.5 mL of 0.9% (w/v) sodium chloride solution. FAMEs were then extracted with 1 mL of hexane. Subsequently, the FAMEs were analyzed on an Agilent 6890N Gas Chromatograph equipped with a 5975 inert XL Mass Selective Detector and Flame Ionization Detector (Agilent Technologies) using a method described in our previous study. ^39,40^ Briefly, FAMEs were separated on a capillary column DB23 (30 222 m×0.25 mm×0.25 µm, Agilent Technologies, Wilmington, DE, USA) using the following program: 2:1 split ratio, 1 µL injection. 4 min at 165°C, then increased to 180°C (10°C/ min) and held for 5 min, and increased to 230°C and held for 5 min.

## 3. Results

### 3.1. Functional validation of pomegranate-derived genes for punicic acid accumulation in yeast biomass

In pomegranate, bifunctional acyl lipid desaturase and conjugase (PgFADX) converts the common fatty acid linoleic acid (LA) on the *sn*-2 position of phosphatidylcholine (PC) to PuA. ^43^ In addition, many biosynthetic pathways upstream and downstream of the PgFADX-catalyzed reaction may synergize effectively with each other for efficient PuA production. ^44,45^ These reactions form the acyl-editing and TAG assembly network (Fig. 1A), where specialized enzymes collaborate to exchange specific acyl groups, redistributing them across various lipid pools and enriching the target fatty acid within the TAG fraction. ^44^ Since most pomegranate enzymes involved in this process have not been fully studied, the identification and isolation of the genes encoding these enzymes were first conducted. As listed in Table S4, genes encoding three pomegranate diacylglycerol acyltransferase 2 (DGAT2), three phospholipid: diacylglycerol acyltransferase (PDAT), and one of each of the following enzymes: diacylglycerol acyltransferase 1 (DGAT1), phosphatidylcholine: diacylglycerol cholinephosphotransferase (PDCT), lysophosphatidylcholine acyltransferase (LPCAT), glycerol-3-phosphate acyltransferase 9 (GPAT9), lysophosphatidic acid acyltransferase 2 (LPAT2), phospholipase A_2_ (PLA_2_), phospholipase C (PLC), and long-chain acyl-CoA synthetase 8 (LACS8) were obtained.

**Fig. 1.**
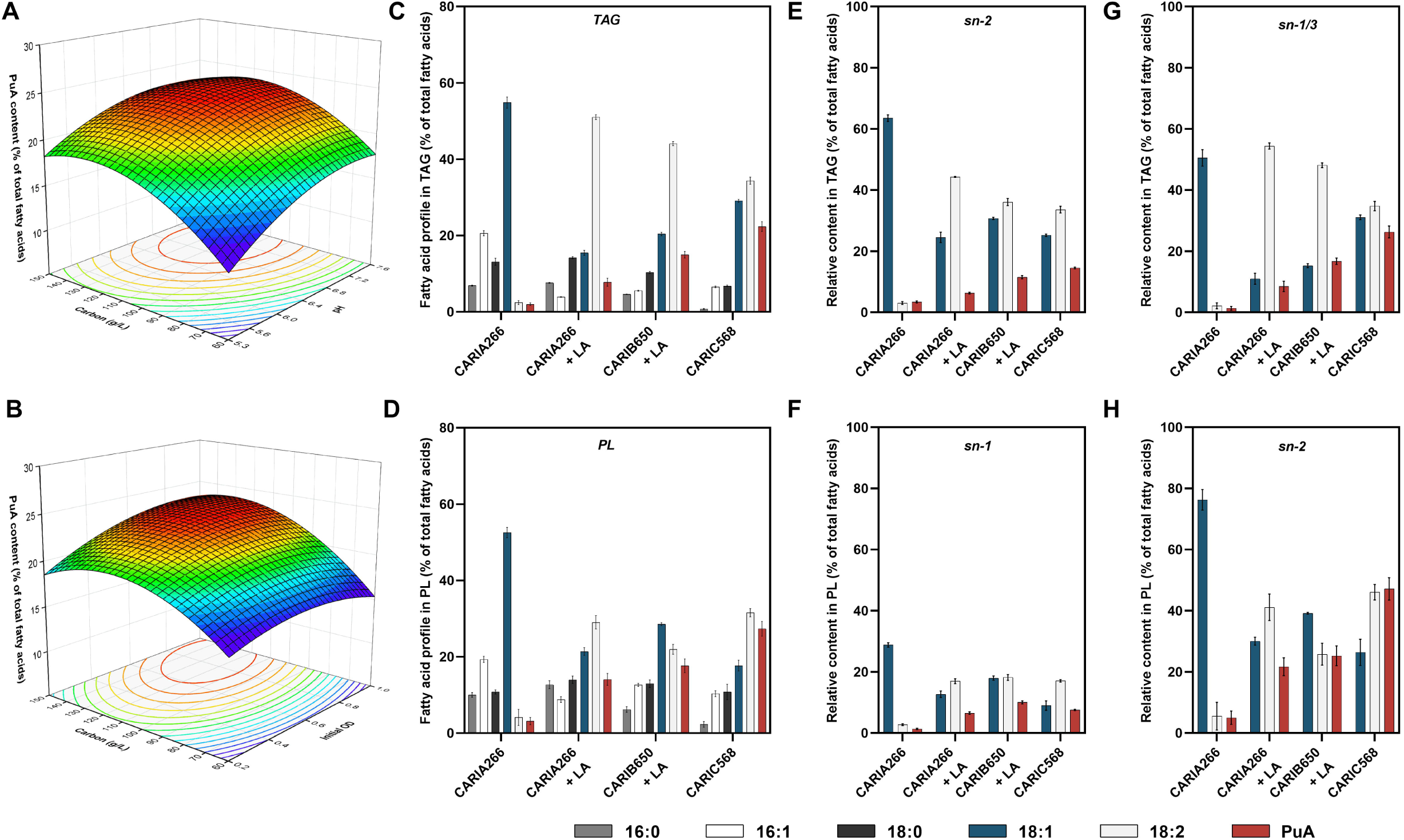
TAG and unusual fatty acid synthetic pathways. (A) Rewiring the lipid biosynthetic pathway in *S. cerevisiae* to produce PuA. Yellow ellipses indicate *S. cerevisiae* native enzymes. Green ellipses indicate enzymes from *P. granatum*. Blue ellipses indicate enzymes from other sources. DGAT/DGA1, acyl-CoA: diacylglycerol acyltransferase; GPAT/SCT1/GPT2, glycerol-3-phosphate acyltransferase; LACS/FAA, long-chain acyl-CoA synthetase; LPAT/SLC1, lysophosphatidic acid acyltransferase; LPC, lysophosphatidylcholine; LPCAT, lysophosphatidylcholine acyltransferase; PAP/APP1/PAH1, phosphatidate phosphatase; PC, phosphatidylcholine; PDAT/LRO1, phospholipid: diacylglycerol acyltransferase; PDCT, phosphatidylcholine: diacylglycerol cholinephosphotransferase; PLA2, phospholipase A2; ACC1, acetyl-CoA carboxylase; FAS, fatty acid synthase; OLE1, acyl-CoA desaturase; ELO, fatty acid elongase; LOA1, lysophosphatidic acid: oleoyl-CoA acyltransferase; DGK1, diacylglycerol kinase; CDS1, phosphatidate cytidylyltransferase; EPT1, choline/ethanolamine phosphotransferase; PIS1, CDP-diacylglycerol--inositol 3-phosphatidyltransferase; PSD, phosphatidylserine decarboxylase; CPT, cholinephosphotransferase; PLC, phospholipase C; ALE1, lysophospholipid acyltransferase; FADX, bifunctional fatty acid conjugase/Delta(12)-oleate desaturase; FAD2, delta(12)-fatty-acid desaturase; PI, phosphatidylinositol; PE, phosphatidylethanolamine; CHO1, CDP-diacylglycerol-serine O-phosphatidyltransferase; PS, phosphatidylserine. Made with biorender.com. (B, C) Functional complementation assay in yeast strain H1246 without/with pomegranate oil hydrolysate. Lane 1-9, pomegranate oil, empty vector control, PgDGAT1, PgDGAT2a, PgDGAT2b, PgDGAT2c, PgPDAT.a, PgPDAT.b, PgPDAT.c respectively. (D) Fatty acid profiles of recombinant H1246 strains expressing different TAG assembly genes in the absence or presence of pomegranate oil hydrolysate. (E) PuA level in yeast strains transformed with various PgFADX protein fusions. Data represents mean ± SD of triplicates.

Since DGAT and PDAT catalyze the crucial final committed step in plant TAG biosynthesis, ^46^ the genes encode PgDGAT1, PgDGAT2.a, PgDGAT2.b, PgDGAT2.c, PgPDAT.a, PgPDAT.b, and PgPDAT.c were transformed into *S. cerevisiae* H1246, a quadruple mutant strain incapable of synthesizing storage lipids including TAG. ^47^ As shown in Fig. 1B, yeast strains expressing *PgDGAT1*, *PgDGAT2.a*, and *PgDGAT2.b*, respectively, restored TAG synthesis ability. Since pomegranate oil contains a high level of PuA, it is plausible to assume that pomegranate-derived TAG assembly enzymes may evolved a preference for PuA-containing substrate. To test this hypothesis, the recombinant yeast strains were fed with pomegranate oil hydrolysate. The results indicated that the strains hosting pomegranate PgDGAT2.c and PgPDAT.a, respectively, could incorporate pomegranate oil hydrolysate to TAG (Fig. 1C). Specifically, the expression of *PgDGAT2.c* led to PuA comprising 20% of the total fatty acids in TAG, suggesting a potential substrate preference for this conjugated linolenic acid (Fig. 1D).

In previous studies, various protein fusions have been used to enhance enzyme performance in metabolic engineering efforts. ^48,49^ In pomegranate, PgFADX is responsible for converting the common fatty acid LA to PuA through a desaturase-like reaction, which requires electron donors, molecular oxygen, and the acyl carrier PC. ^50,51,43^ In addition, H_2_O_2_ may also inhibit the activity of desaturase-like enzymes. ^52^ Therefore, selected genes encoding the PC-binding protein (SCP2), oxygen carrier-protein, soluble domain of the electron transporter cytochrome b5 (CB5SD), and catalase were linked with PgFADX, respectively. The expression cassettes containing these fusions were transformed into BY4741 *snf2*Δ mutant, which exhibited an elevated level of fatty acid synthesis. ^53^ As shown in Fig. 1E, except for the fusion with catalase, all other PgFADX variants showed a slight increase in the PuA level. In addition, yeast SNF1 (sucrose non-fermenting 1) is a protein kinase and master regulator that plays a critical role in regulating yeast energy metabolism and glucose homeostasis, ^54^ and its deletion could lead to enhanced lipid biosynthesis in yeast cells. ^55^ Therefore, we also constructed a BY4741 *snf2*Δ *snf1*Δ double knockout mutant by replacing the *snf1* gene with an *AtSCP2-PgFADX* expression cassette. As shown in Fig. 1E, when *AtCB5SD-PgFADX* was expressed in this strain mutant background, PuA accounted for 6% of the total fatty acids.

### 3.2. Improved bioconversion from linoleic acid to punicic acid by modified yeast strains

The above results were obtained with the plasmid-based gene expression approach, which is less stable without selective pressure and may lead to lower expression levels as well as *in vivo* enzyme concentrations. ^56,57^ Therefore, following these experiments on gene characterization, a workflow was developed for effective integration and testing of PuA accumulation-associated genes directly on the yeast genome. The yeast Ty retrotransposon elements were chosen as the locus for integration, and CRISPR-Cas9 systems were used to facilitate the process. *S. cerevisiae* genome contains a large number of Ty retrotransposon elements. ^58^ Since they do not participate in yeast metabolism, the integration of genes into these regions will lead to higher copy numbers with lower interference with cellular fitness. ^59,60^ Therefore, sequence analysis of yeast Ty retrotransposon and its long terminal repeats (delta sequence) was conducted first. Based on the result, *YERCdelta20* (Table S5), *YDRWdelta23* (Table S6), and *TyA Gag* gene (Table S7) were chosen as the targets for gene shuffling due to their high similarity with other analyzed sequences.

The proposed workflow and designed donor DNA structure are illustrated in Fig. 2A and Fig. 2B, respectively. Various donor DNAs containing expression cassettes were constructed by flanking candidate genes with homology arms targeting yeast Ty retrotransposon. Selection markers with defective promoter regions were also included to enhance the probability of obtaining positive transformations with higher gene copy numbers. Next, the cassettes were co-transformed into yeast cells with the CRISPR-Cas9 system. Various transformants were then individually cultured in small tubes and tested for their intracellular lipid and PuA levels. Based on our previous results, ^61^ 0.03% LA was added to the liquid culture in the first and second round screening as the fatty acid precursor but was omitted in the third-round screening, which aimed for PuA neosynthesis.

**Fig. 2.**
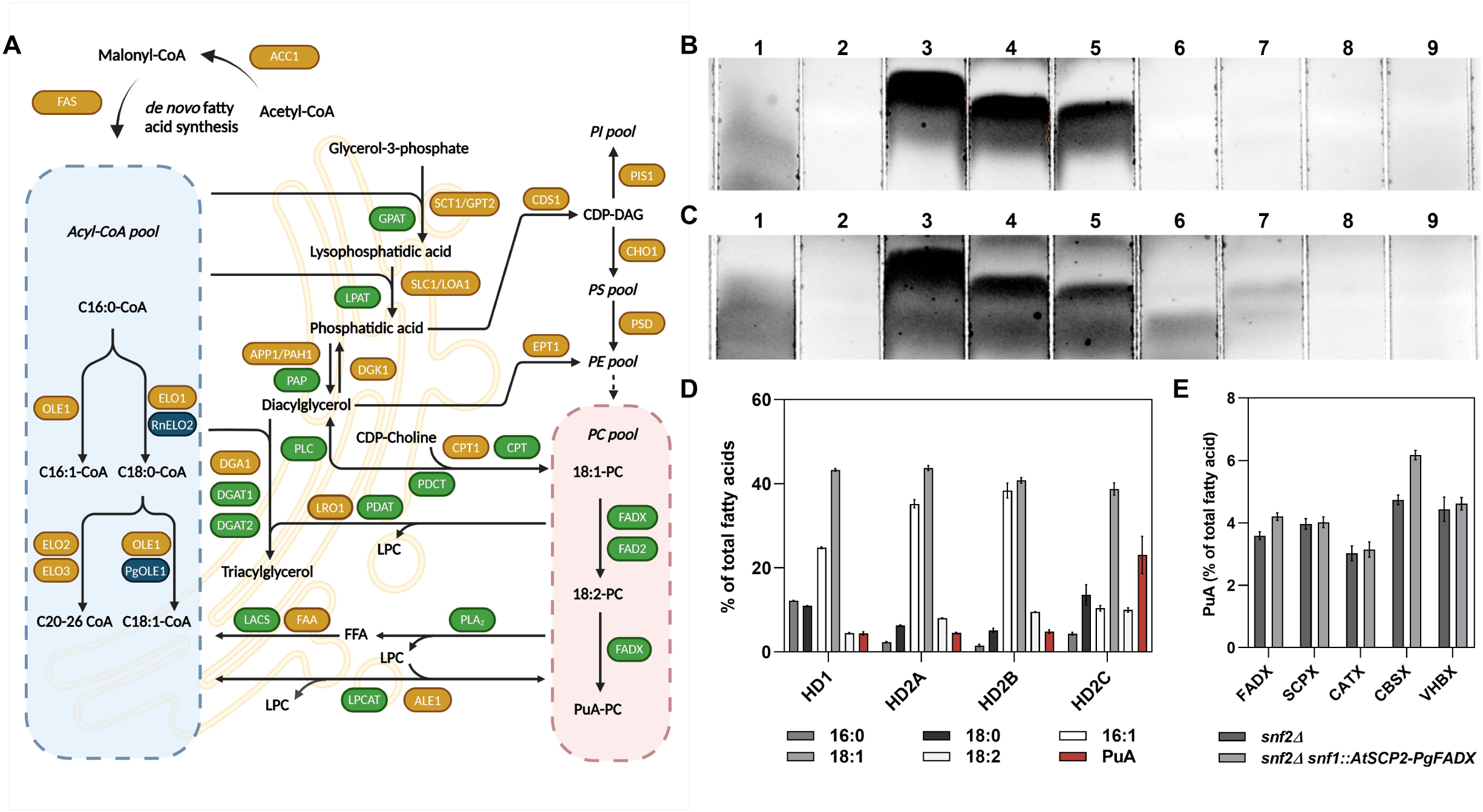
Workflow of the Ty retrotransposon-targeted random gene shuffling. (A) Schematic diagram depicting the screening procedure for Ty retrotransposon-targeted random gene shuffling by CRISPR-Cas9. (B) Schematic diagram of donor DNA design for integration into yeast Ty retrotransposons. Made with biorender.com.

Given the critical role of PgFADX, we integrated the *AtCB5SD-PgFADX* expression cassette into the BY4741 *snf2*Δ *snf1*Δ double knockout mutant during the first round of yeast Ty retrotransposon-targeted integration and subsequently screened 360 strains. With LA feeding, PuA accumulation was detected in all transformants (Fig. 3A). Only a small number of yeast strains (10%) accumulated less than 6% of total fatty acids as PuA. The majority of the strains (84%) accumulated 6%-10% of PuA, and 6% strains accumulated over 10% of PuA. The PuA content in the best strain, designated as CARIA266, reached nearly 11%. To further probe the performance of CARIA266, a growth analysis in shake flasks under different culturing temperatures was conducted. As shown in Fig. 3B, CARIA266 maintained a high PuA level of up to 12.6% at 96 h of cultivation under 20°C. Compared to 30°C, CARIA266 accumulated more monounsaturated fatty acid under 20°C, and the level of PuA gradually increased during the 4-day periods. When cultured under 30°C, the level of PuA was relatively stable and slightly lower than the level obtained under 20°C. The highest titer of PuA was 69 mg/L on day 5.

**Fig. 3.**
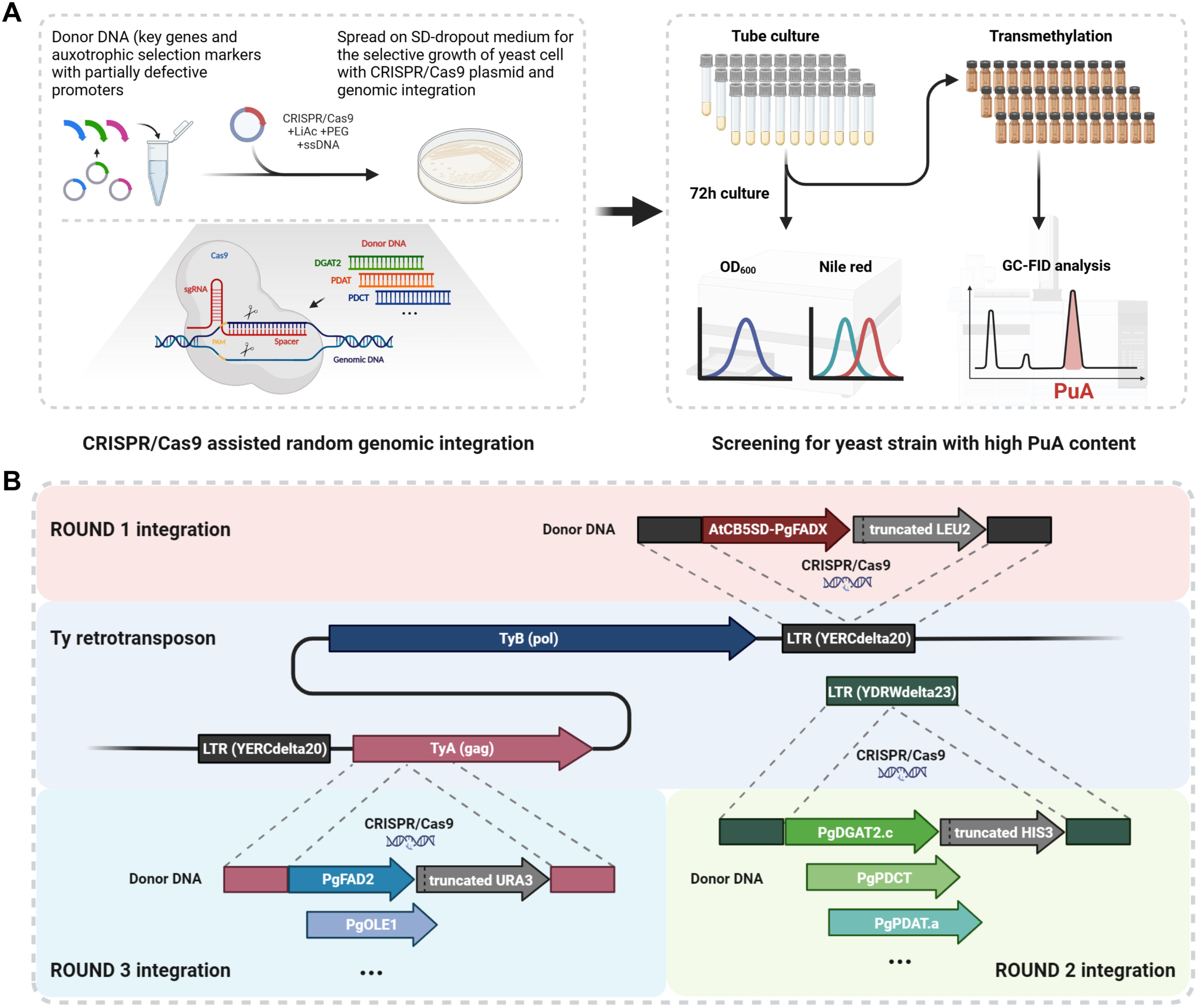
PuA production in *S. cerevisiae* strains constructed by Ty retrotransposon-targeted random gene shuffling. (A, C, E) Three-dimensional scatter plots of the results from round 1, round 2, and round 3 CRISPR-Cas9-assisted random integration, respectively. Each red dot represents an individual yeast transformant. The yeast transformant’s PuA content (represented by the percentage of PuA in total fatty acids, Z-axis), PuA amount (represented by the integrated area of PuA’s peak, Y-axis), and neutral lipid level (represented by the Nile red fluorescence, X-axis) are shown. (B, D, F) Fatty acid composition and PuA production of CARIA266, CARIB650, and CARIC568 at different incubation times under different culturing temperatures. Data represents mean ± SD of triplicates.

Notably, CARIA266 also accumulated a high content of LA, which was not converted to PuA (Fig. 3B). Even though the PuA level in yeast strains obtained by Ty retrotransposon-targeted integration was significantly higher than those generated via the plasmid-based approach, the unexpected buildup of LA indicated that this fatty acid precursor was not efficiently edited, and the native TAG synthesis in yeast may not favor PuA accumulation. One possible cause was the lack of specialized enzymes for acyl-editing and TAG assembly of PuA (Fig. 1A). Therefore, the round two Ty retrotransposon-targeted random gene shuffling focused on the introduction of those enzymes from pomegranate. Since PgDGAT1, PgDGAT2a, PgDGAT2b, PgDGAT2c, and PgPDAT.a could restore TAG synthesis in yeast H1246 (Fig. 1B and 1C), their encoding genes along with *PgPDCT*, *PgLPCAT*, *PgGPAT9*, *PgLPAT2*, *PgPLA_2_*, *PgPLC*, and *PgLACS8* were selected as the candidates.

All these donor DNAs were pooled together and co-transformed into CARIA266 with *YDRWdelta23*-targeting CRISPR-Cas9, and the transformants were cultured in the presence of 0.03% LA. Subsequently, in the 792 screened strains, 70% of them produced 8%-12% of total fatty acids as PuA and 7% accumulated PuA higher than 12% of total fatty acids (Fig. 3C). Compared to the first-round integration (Fig. 3A), the average amount of the PuA from round two strains was higher (Fig. 3C), indicating the enhanced production of PuA from additional gene combination. When the best strain, designated as CARIB650, was cultured in the shake flask under 20°C, it accumulated 18% of total fatty acids as PuA, with the highest PuA titer of 94 mg/L (Fig. 3D). Compared to CARIA266, the content of LA in CARIB650 on day 4 significantly dropped by 25%, indicating an improved conversion rate of this fatty acid precursor.

### 3.3. Developing yeast biomass capable of punicic acid neosynthesis

The above results were obtained by bioconversion of manually fed LA precursor to PuA. Considering the higher costs and the need for consistent quality of precursors required by the bioconversion method, neosynthesis might represent a more advantageous choice. Therefore, the third round of integration was conducted to reconstitute and increase LA precursor supply *in vivo*. Codon-optimized donor DNAs encoding PuA-producing AtCB5SD-PgFADX, LA-producing PgFAD2, oleic acid-producing *Puccinia graminis* acyl-CoA desaturase (PgOLE1), and C18 fatty acid-producing *Rattus norvegicus* fatty acid elongase (RnELO2) (Table S8) were selected as the candidates for integrating into CARIB650. In total, 600 strains were screened without LA feeding. Via neosynthesis, 3% of the strains produced 10%-14% PuA and 0.8% produced PuA higher than 14% of total fatty acids (Fig. 3E). In this round of gene shuffling, 80% of the strains only produced 1%-2% PuA, which was possibly caused by the lack of *PgFAD2* integration. In shake flask cultivation, the best strain (CARIC568) could accumulate 18% PuA (102 mg/L) under 20°C in 4 days without LA feeding (Fig. 3F). Taken together, 1752 yeast transformants were screened in three rounds using the workflow generated in this study. The PuA content and titer have been improved significantly, and the content of total unsaturated fatty acid was increased to 90% in CARIC568 (without LA feeding).

Subsequently, on the level of shake flask culture, a response surface methodology using a multifactorial Box-Behnken design was employed for culture condition optimization to explore the potential of CARIC568 in PuA production. Results revealed carbon source level and initial pH were the most significant factors affecting PuA content and production in CARIC568 (Fig. 4A, 4B, and Fig. S1, S2). Next, second-order polynomial equations were fitted to the experimental results to predict the optimal points (Table S3), which led to optimized conditions including 12% glucose, pH 7.04, and an initial OD_600_ of 0.72. A verification experiment was then carried out to determine the accuracy of the prediction. Following 120 h of incubation in this optimized condition, CARIC568 produced 425 mg/L PuA (Table 1). The lipid content reached 15.5% of dry biomass, with PuA accounting for 26.7% of total fatty acids. Subsequent analysis of genomic DNA revealed that CARIC568 hosted *AtCB5SD-PgFADX* as well as *PgPDCT, PgLPCAT, PgDGAT2.c*, *PgFAD2* and *RnELO2*.

**Fig. 4.**
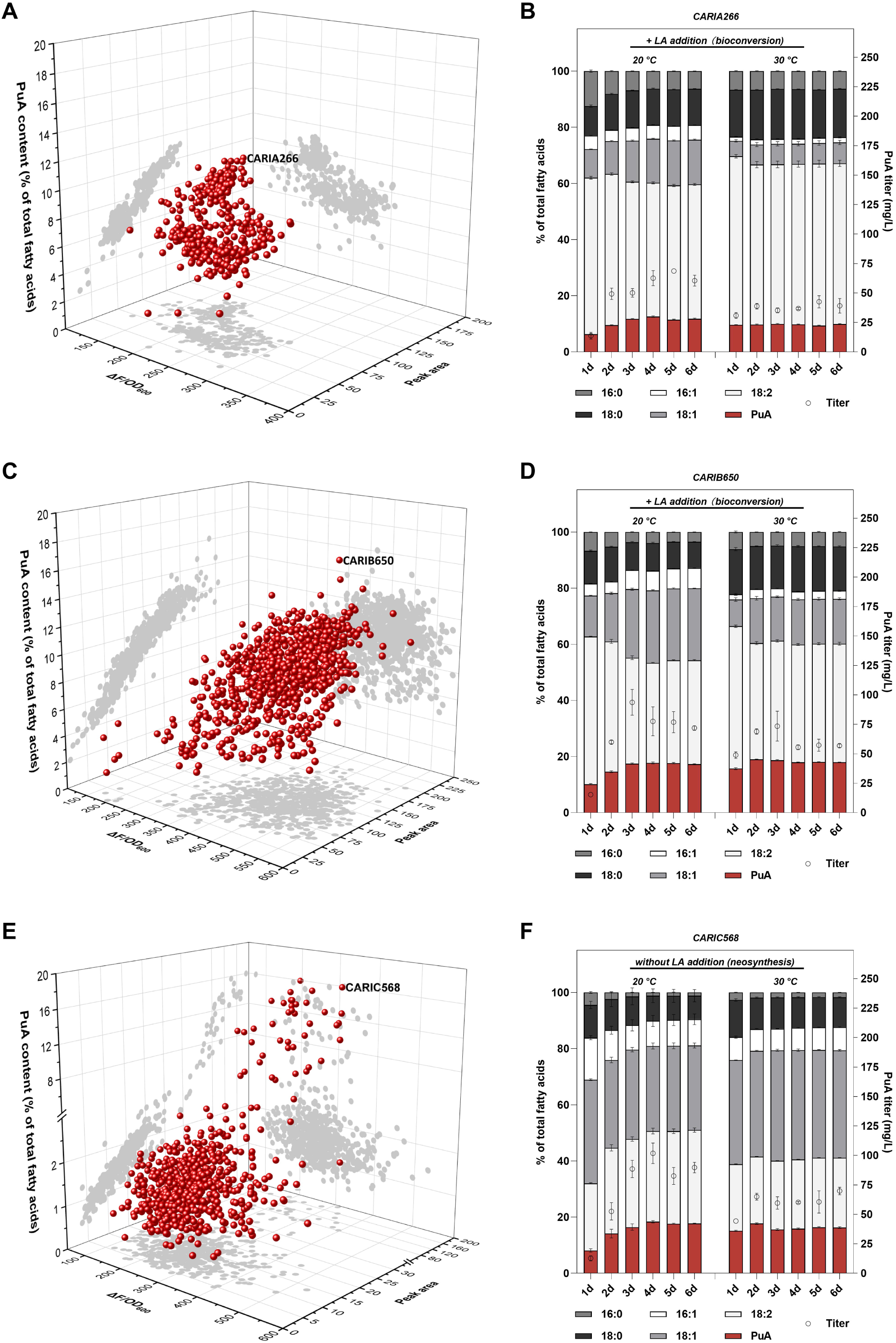
Growth condition optimization and the relative content of PuA in TAG and PL. (A, B) Three-dimensional surface plot of PuA content’s response to carbon source level, and initial pH or initial OD_600_. (C) Fatty acid composition of TAG. (D) Fatty acid composition of PL. (E) Fatty acid composition at the *sn*-2 position of TAG. (F) Fatty acid composition at the *sn*-1 position of PL. (G) The relative content of fatty acid at the *sn*-1/3 positions of TAG. (H) The relative content of fatty acid at the *sn*-2 position of PL. Data represents mean ± SD of triplicates.

**Table 1.**
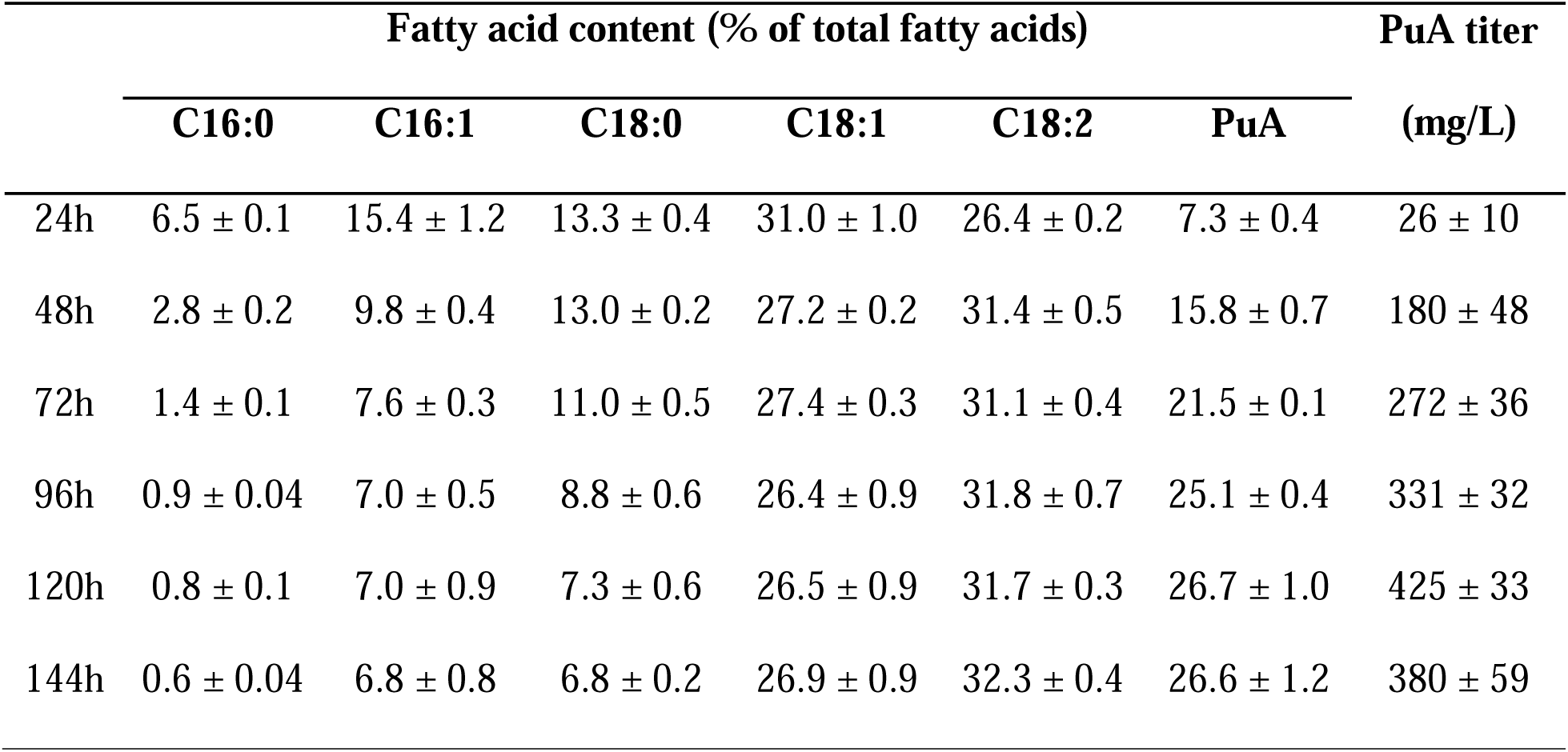
Fatty acid profile and PuA production of CARIC568 over a 6-day growth period in the optimized medium.

### 3.4. Analysis of lipid profile and fatty acid distribution in yeast biomass

To reveal how the lipid profile was influenced by the above engineering efforts, and to assess the impact of varying levels of PuA accumulation, extensive lipid analysis was conducted. This analysis encompassed four groups of samples across three strains, which exhibited low to high amounts of PuA. The conditions tested were as follows: CARIA266 and CARIC568 cultured without LA feeding, and CARIA266 and CARIB650 cultured with 0.03% LA feeding. As shown in Fig. 4C and 4D, when LA was omitted in the culture medium, the PuA level in CARIA266 was relatively low. When LA was available, CARIA266 accumulated 7.8% and 51% of TAG as PuA and LA, respectively. After transforming pomegranate acyl-editing and TAG assembly genes to the genome, CARIB650 accumulated 15% PuA in TAG, representing a 92% increase over CARIA266. Meanwhile, the content of LA in both TAG and PL fractions was lowered in CARIB650. By synthesizing LA directly on PC, CARIC568 accumulated the highest PuA (22.4%) in the TAG fraction compared to other strains.

Subsequently, positional analysis of TAG was conducted, and the results revealed that, in CARIA266 with LA feeding, the *sn*-2 position of TAG contained 44.3% LA and 6.3% PuA (Fig. 4E). After the second round of integration, the LA content of CARIB650 at the *sn*-2 position of TAG was decreased by 19% whereas PuA was increased by 81.7%. In terms of CARIC568, PuA accounted for 14.5% of the fatty acids at the *sn*-2 position of TAG (Fig. 4E) and 26.3% of the fatty acids at the *sn*-1/3 position of TAG (Fig. 4G), suggesting a slight preference for enriching PuA at the *sn*-1/3 position in this strain. Positional analysis of PL indicated that the *sn*-1 position generally contained relatively lower levels of PuA (Fig. 4F). In contrast, PuA accounted for 21.7% (CARIA266+LA), 25.3% (CARIB650+LA), and 47.1% (CARIC568) of the fatty acids at the *sn*-2 position of PL (Fig. 4H), which is consistent with the widely held opinion that the *sn*-2 position of PL is the location for acyl-editing.

## 4. Discussion

PuA is a plant-derived, edible conjugated linolenic acid with various bioactivities. Given its high value, various plants and microorganisms have been engineered to produce this health-promoting fatty acid. Nevertheless, initial efforts to produce PuA in transgenic plants have only led to limited success so far. ^39,40^ For instance, while PuA is the major fatty acid species in pomegranate seed TAG, engineered transgenic *Brassica napus* and *Arabidopsis thaliana* only accumulate 6.6% and 10.6% of PuA in seed TAG fractions, respectively. ^39,40^ Although microbial fermentation presents a promising alternative solution, improving PuA production in microorganisms via metabolic engineering also remains highly challenging. The co-expression of *PgFADX* and *PgFAD2* in *S. cerevisiae* or the single expression of *PgFADX* under the control of a strong promoter in comprehensively modified *Y. lipolytica* only led to less than 1% PuA in yeast single-cell oil. ^11,61^

So far, the exact roles of enzymes involved in PuA synthesis, accumulation, and regulation are yet to be fully elucidated. Individually determining the functions and optimal expression levels of these proteins in plant or microbial hosts adds to the challenge of developing an efficient platform for the heterologous production of PuA. Instead of using traditional gene stacking strategies, where the exact gene candidates, combinations, and approximate ratios were designed before the experiment, this study combined and integrated potentially necessary pathway genes into the yeast genome using CRISPR-Cas9, resulting in a pool of transformants. Subsequent screening of the yeast colonies identified a strain (CARIC568) with high PuA content and a gene combination conducive to PuA accumulation. This method may facilitate the production of other plant-derived unusual fatty acids in microorganisms.

In addition to the results-driven approach designed in this study, it would be interesting to further optimize other aspects for better performance. Although *S. cerevisiae* has been extensively used as a model microorganism to study plant enzymes, some plant-derived enzymes might accumulate poorly in yeast cells. ^37^ This could potentially account for the variability in the results of the TAG synthesis complementation assays conducted in this study (Fig. 1B and 1C). In addition, the stability and turnover rate of PgFADX might also play a role in the process of PuA production. For instance, a previous study showed the N-terminal sequences of plant fatty acid desaturase confer a rapid degradation of the protein. ^62^ Given the structural similarities between various unusual fatty acid-producing enzymes and fatty acid desaturase, further characterization of their N-terminus and protein half-life in transgenic plant and yeast cells could aid the design of a more stable enzyme, thereby improving unusual fatty acid accumulation.

When *S. cerevisiae* was cultured under 30°C, C16 fatty acids accounted for over 60% of its total fatty acid composition. ^63,64^ In comparison, total C16 fatty acid only accounted for 9% at 72 h in CARIC568 grown at 20°C (Table 1). Combining the effects of PgFAD2 and RnELO2, these enzymes greatly contributed to the *in vivo* increase of the C18 fatty acid precursor level and stimulated PuA production. Regarding the genes introduced in the second round of integration, *PDCT*, *LPCAT*, and *DGAT2* are particularly important for enriching unusual fatty acids in plants. A previous study revealed that castor RcPDCT and RcDGAT2 may work in concert to increase tri-ricinoleoyl glycerol levels in castor seeds while also generating nonricinoleate lipids for membrane biosynthesis. ^65^ As illustrated in Fig. 1A, plant LPCAT was also involved in this process by catalyzing both the forward reaction to synthesize PC and the reverse reaction to release acyl-CoA. ^66^ Certain plant LPCAT was shown to have low activities with unusual fatty acid groups in the forward reaction, but higher activity on common fatty acids, ^66,67^ suggesting its role in recruiting specific precursor acyl-CoA into PC for unusual fatty acid synthesis.

It should also be noted that given *S. cerevisiae*’s limited PuA accumulating capabilities observed in our previous study, ^61^ for the second round of integration, we chose to screen pomegranate genes on yeast genome based on the bioconversion method. In the future, it would be interesting to conduct another round of Ty retrotransposon-targeted random gene shuffling of pomegranate acyl-editing and TAG assembly genes on CARIC568, with the approach described in Fig. 2.

Given the presence of a complete pathway for PuA neosynthesis, a better combination of pomegranate genes in terms of *de novo* PuA production would be identified. In addition, considering the potential contribution of yeast native lipid biosynthesis to PuA accumulation, the genes encoding yeast native fatty acid synthesis and TAG assembly enzymes in CARIC568 were kept intact. However, a previous report suggested that isozyme competition might be a limiting factor in the engineering of unusual fatty acid production in heterologous hosts. ^68^ Similarly, in this study, yeast native enzymes may also compete with heterologous enzymes and preferentially integrate fatty acids other than PuA into the glycerol backbone of the TAG. Therefore, replacing them with equivalent heterologous genes in future studies may further enhance PuA levels in yeast lipids.

In summary, genes potentially contributing to plant-derived PuA synthesis were directly integrated and shuffled in the *S. cerevisiae* genome. High throughput screening led to the identification of a recombinant strain capable of accumulating 26.7% of total fatty acids as PuA without LA precursor feeding. In shake flask cultivation, the PuA titer reached 425 mg/L. PuA comprised over 22% of total fatty acids in the TAG fraction of yeast single-cell oil, a significant increase compared to transgenic *A. thaliana* and *B. napus*. With the addition of health-promoting PuA, the engineered yeast biomass may offer enhanced nutritional value over traditional baker’s yeast, making it a potential resource for the food industry.

## CRediT authorship contribution statement

Juli Wang: Conceptualization, Methodology, Investigation, Visualization, Writing—original draft, Writing—review & editing. Guanqun Chen: Conceptualization, Supervision, Funding acquisition, Resources, Writing—review & editing.

### Supporting information

Additional details, including all sequences for the construction of CRISPR-Cas9 and the assembly of pomegranate genes used in this work.

## Acknowledgment

We acknowledge the support provided by the Natural Sciences and Engineering Research Council of Canada (NSERC) Discovery Grant (G.C.), NSERC Alliance grant (ALLRP 561166 – 20), Canada Research Chairs Program (G.C.), Alberta Innovates (2019F111R), Canadian Poultry Research Council (AMN122), Cargill/Diamond V (2019F111R), Results Driven Agriculture Research (2019F111R), Canada Foundation for Innovation-John R. Evans Leaders Fund (Project number 41867) and Research Capacity Program of Alberta (RCP-22-023-SEG). We also acknowledge the financial support to the student including the Alberta Innovates Graduate Student Scholarship (J.W.)

