## Supplementary information for "Engineering *Saccharomyces cerevisiae* for the production of punicic acid-rich yeast biomass"

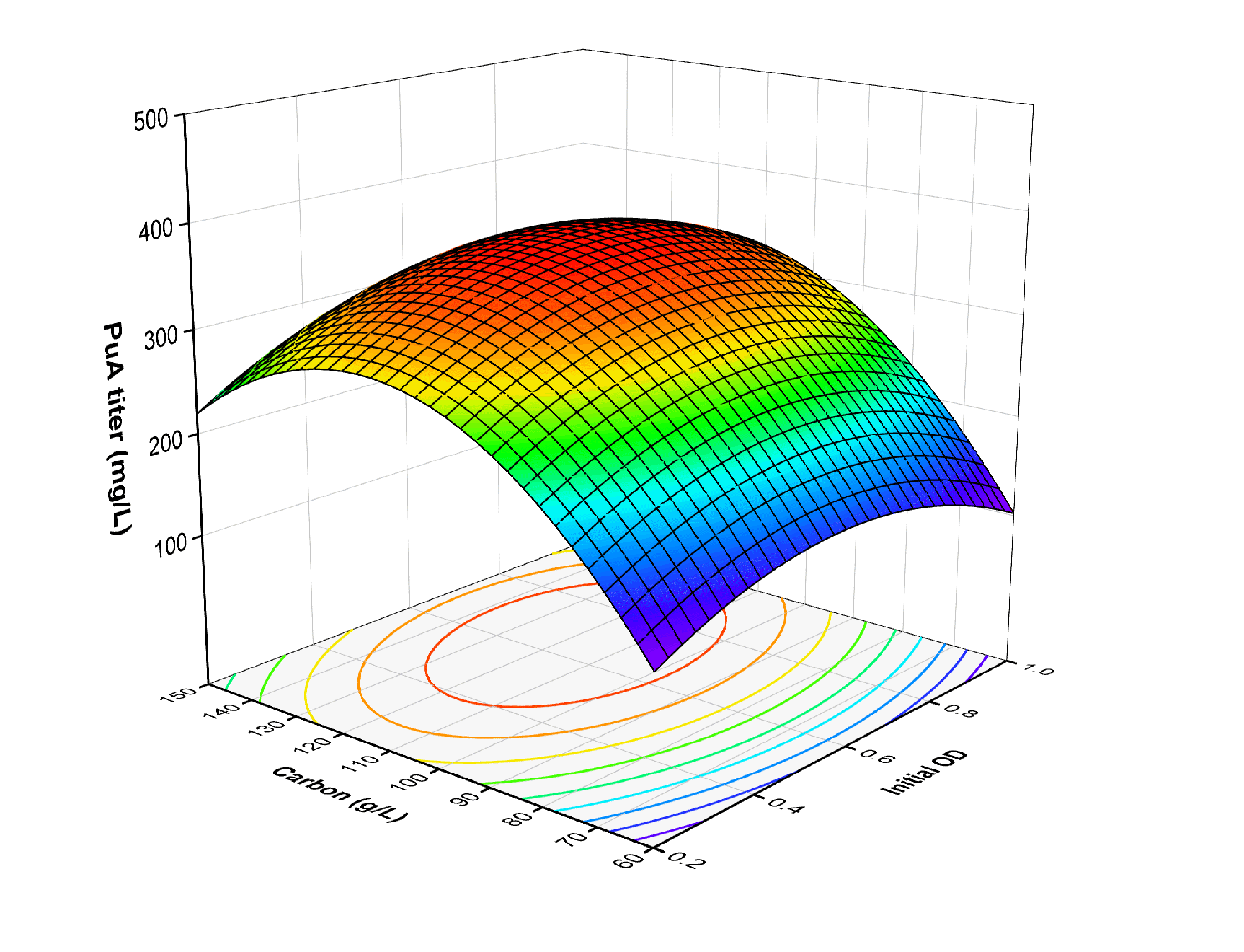

**Fig. S1**. Three-dimensional surface plot of PuA titer to carbon source level and initial OD.

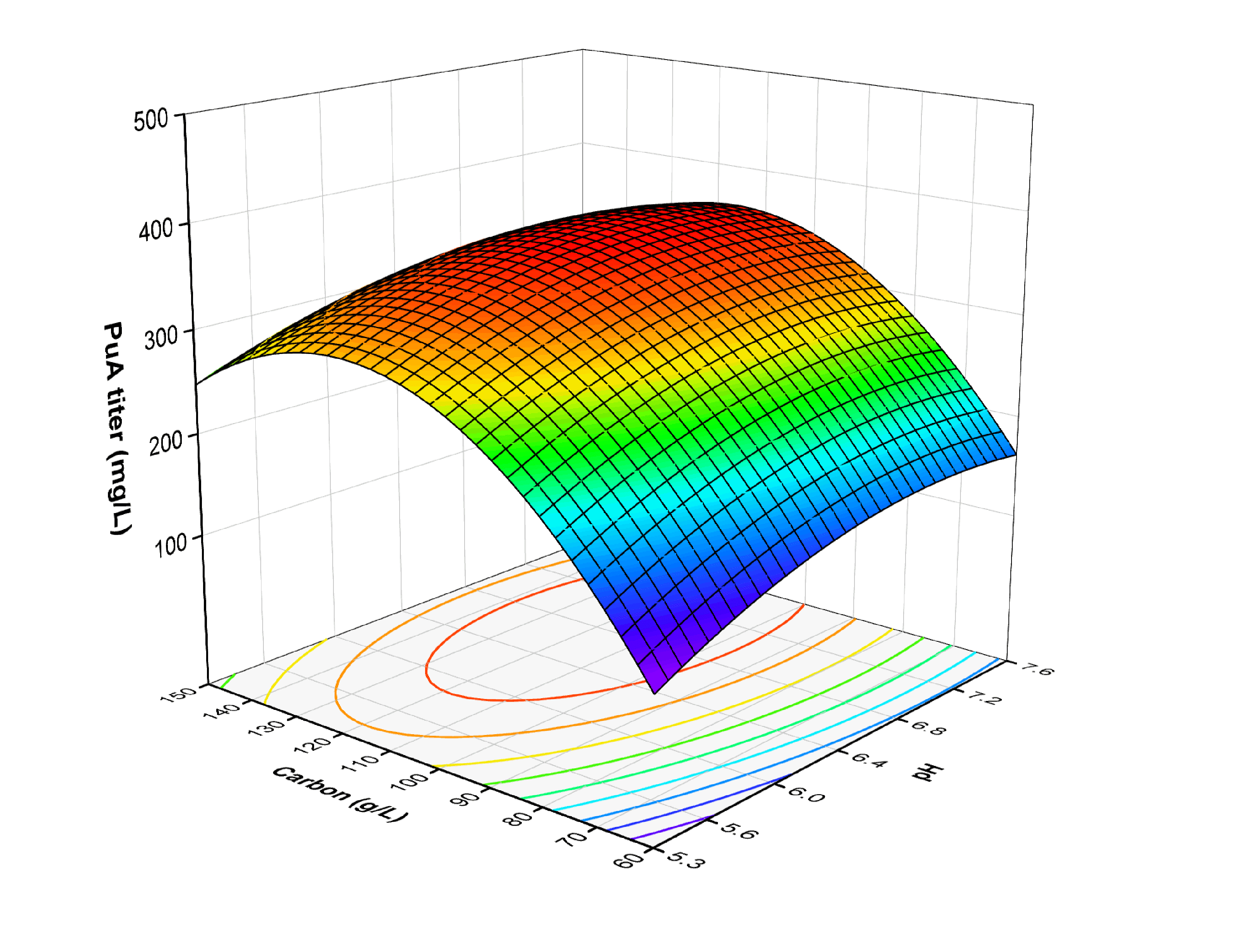

**Fig. S2.** Three-dimensional surface plot of PuA titer to carbon source level and initial pH.

**Table S1.** Strains and plasmids used in this study.

| **Name** | **Description** | **Source** |
| --- | --- | --- |
| **Strains** | | |
| *E. coli* DH5a | F– endA1 glnV44 thi-1 recA1 relA1 gyrA96 deoR nupG purB20 φ80dlacZΔM15Δ(lacZYA-argF) U169, hsdR17(rK–mK+), λ– | Invitrogen |
| *S. cerevisiae* H1246 | *MATα are1Δ::HIS3, are2Δ::LEU2, dga1Δ::KanMX4, lro1Δ::TRP1 ADE2* | Laboratory stock |
| *S. cerevisiae* BY4741 *snf2Δ* | *S. cerevisiae* BY4741-*snf2Δ::KanMX* | Euroscarf |
| *S. cerevisiae* BY4741 *snf2Δ snf1Δ* | *S. cerevisiae* BY4741-*snf2Δ::KanMX-snf1::AtSCP2-PgFADX* | This study |
| HD1 | *S. cerevisiae* H1246/pST3-PgDGAT1 | This study |
| HD2A | *S. cerevisiae* H1246/pST3-PgDGAT2.a | This study |
| HD2B | *S. cerevisiae* H1246/pST3-PgDGAT2.b | This study |
| HD2C | *S. cerevisiae* H1246/pST3-PgDGAT2.c | This study |
| HPA | *S. cerevisiae* H1246/pST3-PgPDAT.a | This study |
| HPB | *S. cerevisiae* H1246/pST3-PgPDAT.b | This study |
| HPC | *S. cerevisiae* H1246/pST3-PgPDAT.c | This study |
| FADX | *S. cerevisiae* *snf2Δ/*pHRPT-PgFADX | This study |
| SCPX-*snf2Δ* | *S. cerevisiae* *snf2Δ/*pHRPT-AtSCP2-PgFADX | This study |
| CATX-*snf2Δ* | *S. cerevisiae* *snf2Δ/*pHRPT-ScCAT-PgFADX | This study |
| CBSX-*snf2Δ* | *S. cerevisiae* *snf2Δ/*pHRPT-AtCB5SD-PgFADX | This study |
| VHBX-*snf2Δ* | *S. cerevisiae* *snf2Δ/*pHRPT-Vhb-PgFADX | This study |
| FADX-*snf1Δsnf2Δ* | *S. cerevisiae* *snf2Δ snf1Δ/*pHRPT-PgFADX | This study |
| SCPX-*snf1Δsnf2Δ* | *S. cerevisiae* *snf2Δ snf1Δ/*pHRPT-AtSCP2-PgFADX | This study |
| CATX-*snf1Δsnf2Δ* | *S. cerevisiae* *snf2Δ snf1Δ/*pHRPT-ScCAT-PgFADX | This study |
| CBSX-*snf1Δsnf2Δ* | *S. cerevisiae* *snf2Δ snf1Δ/*pHRPT-AtCB5SD-PgFADX | This study |
| VHBX-*snf1Δsnf2Δ* | *S. cerevisiae* *snf2Δ snf1Δ/*pHRPT-Vhb-PgFADX | This study |
| CARIC568 | *S. cerevisiae snf2Δ snf1Δ P_PGK1_-AtCB5SD-PgFADX-T_ADH1_-LEU2 P_TEF1_- PgDGAT2.c-T_CYC1_-HIS3 P_TEF1_-PgLPCAT-T_CYC1_-HIS3 P_TEF1_-PgPDCT-T_CYC1_-HIS3 P_GAP_-PgFAD2-T_ADH2_-URA3 P_GAP_-RnELO2-T_ADH2_-URA3* | This study |
| **Plasmids** | | |
| pCRCT | TracrRNA/ iCas9/ Amp^R^/ URA3 | (*33*) |
| pCRCT-MET | TracrRNA/ iCas9/ Amp^R^/ MET15 | This study |
| pDA-CRISPR | pCRCT/ YERCdelta20 crRNA | This study |
| pDB-CRISPR | pCRCT/ YDRWdelta23 crRNA | This study |
| pDC-CRISPR | pCRCT-MET/ TyA Gag gene crRNA | This study |
| pHRPT-AtCB5SD-PgFADX | Amp^R^/URA3/ Hyg^R^/*P_PGK1_-AtCB5SD-PgFADX-T_ADH1_* | This study |
| pHRPT-AtSCP2-PgFADX | Amp^R^/URA3/ Hyg^R^/*P_PGK1_-* AtSCP2-PgFADX*-T_ADH1_* | This study |
| pHRPT-PgFADX | Amp^R^/URA3/ Hyg^R^/*P_PGK1_*-PgFADX *-T_ADH1_* | This study |
| pHRPT-ScCAT-PgFADX | Amp^R^/URA3/ Hyg^R^/*P_PGK1_-*ScCAT-PgFADX*-T_ADH1_* | This study |
| pHRPT-VHB-PgFADX | Amp^R^/URA3/ Hyg^R^/*P_PGK1_*-VHB-PgFADX*-T_ADH1_* | This study |
| pST3-PgDGAT1 | Amp^R^/URA3/ Hyg^R^/*P_TEF1_-*PgDGAT1*-T_CYC1_* | This study |
| pST3-PgDGAT2.a | Amp^R^/URA3/ Hyg^R^/*P_TEF1_-*PgDGAT2.a*-T_CYC1_* | This study |
| pST3-PgDGAT2.b | Amp^R^/URA3/ Hyg^R^/*P_TEF1_-*PgDGAT2.b*-T_CYC1_* | This study |
| pST3-PgDGAT2.c | Amp^R^/URA3/ Hyg^R^/*P_TEF1_-*PgDGAT2.c*-T_CYC1_* | This study |
| pST3-PgGPAT9 | Amp^R^/URA3/ Hyg^R^/*P_TEF1_-*PgGPAT9*-T_CYC1_* | This study |
| pST3-PgLACS8 | Amp^R^/URA3/ Hyg^R^/*P_TEF1_-*PgLACS8*-T_CYC1_* | This study |
| pST3-PgLACS9 | Amp^R^/URA3/ Hyg^R^/*P_TEF1_-*PgLACS9*-T_CYC1_* | This study |
| pST3-PgLPAT2 | Amp^R^/URA3/ Hyg^R^/*P_TEF1_-*PgLPAT2*-T_CYC1_* | This study |
| pST3-PgLPCAT | Amp^R^/URA3/ Hyg^R^/*P_TEF1_-*PgLPCAT*-T_CYC1_* | This study |
| pST3-PgPDAT.a | Amp^R^/URA3/ Hyg^R^/*P_TEF1_-*PgPDAT.a*-T_CYC1_* | This study |
| pST3-PgPDAT.b | Amp^R^/URA3/ Hyg^R^/*P_TEF1_-*PgPDAT.b*-T_CYC1_* | This study |
| pST3-PgPDAT.c | Amp^R^/URA3/ Hyg^R^/*P_TEF1_-*PgPDAT.c*-T_CYC1_* | This study |
| pST3-PgPDCT | Amp^R^/URA3/ Hyg^R^/*P_TEF1_-*PgPDCT*-T_CYC1_* | This study |
| pST3-PgPLA2a | Amp^R^/URA3/ Hyg^R^/*P_TEF1_-*PgPLA2a*-T_CYC1_* | This study |
| pST3-PgPLC | Amp^R^/URA3/ Hyg^R^/*P_TEF1_-*PgPLC*-T_CYC1_* | This study |
| pDAL-AtCB5SD-PgFADX | *YERCdelta20-left arm/P_PGK1_-AtCB5SD-PgFADX-T_ADH1_-LEU2/YERCdelta20-right arm* | This study |
| pDBH-PgDGAT1 | *YDRWdelta23-left arm/P_TEF1_-PgDGAT1-T_CYC1_/YDRWdelta23-right arm* | This study |
| pDBH-PgDGAT2.a | *YDRWdelta23-left arm/P_TEF1_-PgDGAT2.a-T_CYC1_/YDRWdelta23-right arm* | This study |
| pDBH-PgDGAT2.b | *YDRWdelta23-left arm/P_TEF1_-PgDGAT2.b-T_CYC1_/YDRWdelta23-right arm* | This study |
| pDBH-PgDGAT2.c | *YDRWdelta23-left arm/P_TEF1_-PgDGAT2.c-T_CYC1_/YDRWdelta23-right arm* | This study |
| pDBH-PgGPAT9 | *YDRWdelta23-left arm/P_TEF1_-PgGPAT9-T_CYC1_/YDRWdelta23-right arm* | This study |
| pDBH-PgLACS8 | *YDRWdelta23-left arm/P_TEF1_-PgLACS8-T_CYC1_/YDRWdelta23-right arm* | This study |
| pDBH-PgLPAT2 | *YDRWdelta23-left arm/P_TEF1_-PgLPAT2-T_CYC1_/YDRWdelta23-right arm* | This study |
| pDBH-PgLPCAT | *YDRWdelta23-left arm/P_TEF1_-PgLPCAT-T_CYC1_/YDRWdelta23-right arm* | This study |
| pDBH-PgPDAT.a | *YDRWdelta23-left arm/P_TEF1_-PgPDAT.a-T_CYC1_/YDRWdelta23-right arm* | This study |
| pDBH-PgPDCT | *YDRWdelta23-left arm/P_TEF1_-PgPDCT-T_CYC1_/YDRWdelta23-right arm* | This study |
| pDBH-PgPLA2a | *YDRWdelta23-left arm/P_TEF1_-PgPLA2a-T_CYC1_/YDRWdelta23-right arm* | This study |
| pDBH-PgPLC | *YDRWdelta23-left arm/P_TEF1_-PgPLC-T_CYC1_/YDRWdelta23-right arm* | This study |
| pDCU-AtCB5SD-FADX OPED | *TyA Gag-left arm/P_GAP_-AtCB5SD-FADX OPED-T_ADH2_-URA3/ TyA Gag-right arm* | This study |
| pDCU-PgFAD2 OPED | *TyA Gag-left arm/P_GAP_-PgFAD2 OPED-T_ADH2_-URA3/ TyA Gag-right arm* | This study |
| pDCU-PgOLE1 OPED | *TyA Gag-left arm/P_GAP_-PgOLE1 OPED-T_ADH2_-URA3/ TyA Gag-right arm* | This study |
| pDCU-RnELO2 OPED | *TyA Gag-left arm/P_GAP_-RnELO2 OPED-T_ADH2_-URA3/ TyA Gag-right arm* | This study |

**Table S2.** Guide sequences and target region used in this study.

| **Target region** | **Sequence** |
| --- | --- |
| *crRNAYERCdelta20* | TATACTAGAAGTTCTCCTCG/**AGG** |
| *crRNAYDRWdelta23* | GCAAGGATTGATAATGTAAT/**AGG** |
| *crRNATyAGag* | TCAGGTGATGGAGTGCTCAG/**AGG** |
| *YERCdelta20* | TGTTGGAATAGAAATCAACTATCATCTACTAACTAGTATTTACATTACTAGTATATTATCATATACGGTGTTAGAAGATGACGCAAATGATGAGAAATAGTCATCTAAATTAGTGGAAGCTGAAACGCAAGGATTGATAATGTAATAGGATCAATGAATATAAACATATAAAACGGAATGAGGAATAATCGTAATATTAGTATGTAGAAATATAGATTCCATTTTGAGGATTCCTATATCCTCGAGGAGAACTTCTAGTATATTCTGTATACCTAATATTATAGCCTTTATCAACAATGGAATCCCAACAATTATCTCAACATTCACCCATTTCTCATGGTAGCGCCTGTGCTTCGGTTACTTCTAAGGAAGTCCACACAAATCAAGATCCGTTAGACGTTTCAGCTTCCA |
| *YDRWdelta23* | TGTTGGAATAAAAATCCACTATCGTCTATCAACTAATAGTTATATTATCAATATATTATCATATACGGTGTTAAGATGATGACATAAGTTATGAGAAGCTGTCATCGATGTTAGAGGAAGCTGAAACGCAAGGATTGATAATGTAATAGGATCAATGAATATAAACATATAAAACGGAATGAGGAATAATCGTAATATTAGTATGTAGAAATATAGATTCCATTTTGAGGATTCCTATATCCTCGA |
| *TyA Gag* | ATGGAATCCCAACAATTATCTCAACATTCACCCATTTCTCATGGTAGCGCCTGTGCTTCGGTTACTTCTAAGGAAGTCCACACAAATCAAGATCCGTTAGACGTTTCAGCTTCCAAAACAGAAGAATGTGAGAAGGCTTCCACTAAGGCTAACTCTCAACAGACAACAACACCTGCTTCATCAGCTGTTCCAGAGAACCCCCATCATGCCTCTCCTCAACCTGCTTCAGTACCACCTCCACAGAATGGGCCGTACCCACAGCAGTGCATGATGACCCAAAACCAAGCCAATCCATCTGGTTGGTCATTTTACGGACACCCATCTATGATTCCGTATACACCTTATCAAATGTCGCCTATGTACTTTCCACCTGGGCCACAATCACAGTTTCCGCAGTATCCATCATCAGTTGGAACGCCTCTGAGCACTCCATCACCTGAGTCAGGTAATACATTTACTGATTCATCCTCAGCGGACTCTGATATGACATCCACTAAAAAATATGTCAGACCACCACCAATGTTAACCTCACCTAATGACTTTCCAAATTGGGTTAAAACATACATCAAATTTTTACAAAACTCGAATCTCGGTGGTATTATTCCGACAGTAAACGGAAAACCCGTACGTCAGATCACTGATGATGAACTCACCTTCTTGTATAACACTTTTCAAATATTTGCTCCCTCTCAATTCCTACCTACCTGGGTCAAAGACATCCTATCCGTTGATTATACGGATATCATGAAAATTCTTTCCAAAAGTATTGAAAAAATGCAATCTGATACCCAAGAGGCAAACGACATTGTGACCCTGGCAAATTTGCAATATAATGGCAGTACACCTGCAGATGCATTTGAAACAAAAGTCACAAACATTATCGACAGACTGAACAATAATGGCATTCATATCAATAACAAGGTCGCATGCCAATTAATTATGAGAGGTCTATCTGGCGAATATAAATTTTTACGCTACACACGTCATCGACATCTAAATATGACAGTCGCTGAACTGTTCTTAGATATCCATGCTATTTATGAAGAACAACAGGGATCGAGAAACAGCAAACCTAATTACAGGAGAAATCTGAGTGATGAGAAGAATGATTCTCGCAGCTATACGAATACAACCAAACCCAAAGTTATAGCTCGGAATCCTCAAAAAACAAATAATTCGAAATCGAAAACAGCCAGGGCTCACAATGTATCCACATCTAATAACTCTCCCAGCACGGACAACGATTCCATCAGTAAATCAACTACTGAACCGATTCAATTGAACAATAAGCACGACCTTCACCTTAGGCCAGGAACTTACTGA |

**Table S3.** Optimization of culture condition using response surface methodology.

| **Run** | **Carbon** | **pH** | **OD** | **Titer mean** | **Content mean** |
| --- | --- | --- | --- | --- | --- |
| 1 | 60 | 6.5 | 0.2 | 102.5 | 15.3 |
| 2 | 105 | 5.3 | 1 | 214.5 | 18.7 |
| 3 | 105 | 6.5 | 0.6 | 330.3 | 23.3 |
| 4 | 105 | 7.6 | 0.2 | 292.8 | 21.2 |
| 5 | 150 | 6.5 | 1 | 278.8 | 23.1 |
| 6 | 150 | 5.3 | 0.6 | 266.5 | 18.2 |
| 7 | 150 | 6.5 | 0.2 | 219.6 | 19.5 |
| 8 | 105 | 6.5 | 0.6 | 331.5 | 21.7 |
| 9 | 60 | 7.6 | 0.6 | 142.6 | 17.7 |
| 10 | 105 | 5.3 | 0.2 | 220.6 | 13.9 |
| 11 | 150 | 7.6 | 0.6 | 265.8 | 21.5 |
| 12 | 105 | 6.5 | 0.6 | 371.7 | 24.4 |
| 13 | 60 | 5.3 | 0.6 | 108.9 | 12.6 |
| 14 | 105 | 6.5 | 0.6 | 380.6 | 22.1 |
| 15 | 105 | 6.5 | 0.6 | 371.4 | 23.2 |
| 16 | 105 | 7.6 | 1 | 343.4 | 23.4 |
| 17 | 60 | 6.5 | 1 | 102.1 | 14.4 |
| PuA content=  22.9458 + 2.78723 * X1 + 2.49504 * X2 + 1.20784 * X3 + -0.461944 * X1X2 + 1.16779 * X1X3 -0.689039 * X2X3 -3.32149 * X1^2 -2.19481 * X2^2 -1.54413 * X3^2 | | | | | |
| PuA titer=  357.115 + 71.6532 * X1 + 27.4837 * X2 + 13.2087 * X3 -8.89518 * X1X2 + 14.9048 * X1X3 + 14.8339 * X2X3 -126.632 * X1^2 -36.2917 * X2^2 -54.7395 * X3^2 | | | | | |

**Table S4.** Pomegranate acyl-editing and TAG assembly genes.

| **Name** | **Query species** | **Query protein** | **Annotation** | **Designation** | **Subject protein** | **Subject CDS** |
| --- | --- | --- | --- | --- | --- | --- |
| *DGAT1* | *Vernicia fordii* | >ABC94472.1 | Tung DGAT1, natural producer of UFA; producer of conjugated 18:3 PUFA | PgDGAT1 | >XP_031382678.1 | >XM_031526818.1:200-1825 |
| *DGAT2* | *Vernicia fordii* | >ABC94474.1 | Tung DGAT2, natural producer of UFA; producer of conjugated 18:3 PUFA | PgDGAT2.a | >XP_031388774.1 | >XM_031532914.1:353-1372 |
|  |  |  |  | PgDGAT2.b | >XP_031401853.1 | >XM_031545993.1:1181-2185 |
|  |  |  |  | PgDGAT2.c | >XP_031401854.1 | >XM_031545994.1:161-1150 |
| *PDAT* | *Linum usitatissimum* | >AHA57448.1 | Flax LuPDAT2, natural producer of 18:3 PUFA, express in vegetative tissue but high protein accmulation in yeast, Similar preference with flax LuPDAT1, which is a seed specific PDAT | PgPDAT.a | >XP_031388206.1 | >XM_031532346.1:1-2055 |
|  |  |  |  | PgPDAT.b | >XP_031380888.1 | >XM_031525028.1:129-2180 |
|  |  |  |  | PgPDAT.c | >XP_031388207.1 | >XM_031532347.1:620-2203 |
| *GPAT9* | *Ricinus communis* | >ACB30546.1 | Castor GPAT9, natural producer of UFA | PgGPAT9 | >XP_031383927.1 | >XM_031528067.1:20-1222 |
| *LPAT2* | *Vernicia fordii* | >AZZ09512.1 | Tung LPAT2, natural producer of UFA; producer of conjugated 18:3 PUFA | PgLPAT2 | >XP_031398075.1 | >XM_031542215.1:232-1395 |
| *LACS8* | *Linum usitatissimum* | >Lus10016083.g | Flax LACS8, natural producer of 18:3 PUFA, has high preference towards18:3 fatty acid | PgLACS8 | >XP_031372442.1 | >XM_031516582.1:362-2551 |
| *LPCAT* | *Linum usitatissimum* | >Lus10006325.g | Flax LPCAT, natural producer of 18:3 PUFA | PgLPCAT | >XP_031394089.1 | >XM_031538229.1:138-1535 |
| *PDCT* | *Linum usitatissimum* | >AHE80679.1 | Flax PDCT, natural producer of 18:3 PUFA | PgPDCT | >XP_031389447.1 | >XM_031533587.1:349-1212 |
| *PLA2a* | *Ricinus communis* | >29840.t000027 | Castor PLA2a, natural producer of UFA | PgPLA2a | >XP_031374260.1 | >XM_031518400.1:165-641 |
| *PLC* | *Ricinus communis* | >30115.t000070 | Castor PLC, natural producer of UFA | PgPLC | >XP_031386572.1 | >XM_031530712.1:225-1511 |

**Table S5.** Sequence similarity and identity search for *YERCdelta20*.

| **Name** | **Max Score** | **Total Score** | **Cover** | **E value** | **Per. ident** | **Acc. Len** |
| --- | --- | --- | --- | --- | --- | --- |
| YPLWdelta4 | 623 | 623 | 100% | 0 | 100 | 337 |
| YPLWdelta3 | 623 | 623 | 100% | 0 | 100 | 337 |
| YCLWdelta15 | 623 | 623 | 100% | 0 | 100 | 337 |
| YARCdelta5 | 623 | 623 | 100% | 0 | 100 | 337 |
| YARCdelta4 | 623 | 623 | 100% | 0 | 100 | 337 |
| YLRCdelta8 | 619 | 619 | 99% | 7.00E-180 | 100 | 335 |
| YLRCdelta7 | 619 | 619 | 99% | 7.00E-180 | 100 | 335 |
| YMLWdelta5 | 612 | 612 | 100% | 1.00E-177 | 99 | 337 |
| YERCdelta19 | 606 | 606 | 100% | 6.00E-176 | 99 | 337 |
| YMLWdelta6 | 562 | 562 | 93% | 1.00E-162 | 99 | 316 |
| YPRCdelta19 | 508 | 508 | 100% | 2.00E-146 | 94 | 334 |
| YPRCdelta18 | 508 | 508 | 100% | 2.00E-146 | 94 | 334 |
| YLRWdelta11 | 508 | 508 | 100% | 2.00E-146 | 94 | 334 |
| YDRWdelta27 | 508 | 508 | 100% | 2.00E-146 | 94 | 334 |
| YNLWdelta4 | 503 | 503 | 100% | 8.00E-145 | 94 | 334 |
| YNLWdelta3 | 503 | 503 | 100% | 8.00E-145 | 94 | 334 |
| YLRWdelta10 | 497 | 497 | 100% | 4.00E-143 | 93 | 334 |
| YJRWdelta12 | 497 | 497 | 100% | 4.00E-143 | 93 | 334 |
| YGRCdelta16 | 497 | 497 | 100% | 4.00E-143 | 93 | 334 |
| YDRWdelta28 | 497 | 497 | 100% | 4.00E-143 | 93 | 334 |
| YDRCdelta22 | 497 | 497 | 100% | 4.00E-143 | 93 | 334 |
| YPRCdelta23 | 490 | 490 | 100% | 6.00E-141 | 93 | 338 |
| YOLWdelta5 | 490 | 490 | 100% | 6.00E-141 | 93 | 338 |
| YLRWdelta13 | 486 | 486 | 100% | 8.00E-140 | 93 | 334 |
| YJRWdelta13 | 484 | 484 | 100% | 3.00E-139 | 93 | 338 |
| YJRWdelta11 | 484 | 484 | 100% | 3.00E-139 | 93 | 338 |
| YGRWdelta14 | 484 | 484 | 100% | 3.00E-139 | 93 | 338 |
| YGRWdelta13 | 484 | 484 | 100% | 3.00E-139 | 93 | 338 |
| YGRCdelta15 | 477 | 477 | 94% | 5.00E-137 | 94 | 334 |
| YPRCdelta24 | 473 | 473 | 100% | 6.00E-136 | 92 | 338 |
| YLRWdelta14 | 466 | 466 | 94% | 1.00E-133 | 93 | 334 |
| YDRCdelta8 | 466 | 466 | 94% | 1.00E-133 | 93 | 334 |
| YDRCdelta7 | 466 | 466 | 94% | 1.00E-133 | 93 | 334 |
| YDRCdelta21 | 466 | 466 | 94% | 1.00E-133 | 93 | 334 |
| YPRWdelta21 | 453 | 453 | 94% | 8.00E-130 | 92 | 338 |
| YPRWdelta20 | 453 | 453 | 94% | 8.00E-130 | 92 | 338 |
| YOLWdelta4 | 453 | 453 | 94% | 8.00E-130 | 92 | 338 |
| YMRCdelta9 | 453 | 453 | 94% | 8.00E-130 | 92 | 338 |
| YMRCdelta10 | 453 | 453 | 94% | 8.00E-130 | 92 | 338 |
| YLRCdelta2 | 453 | 453 | 94% | 8.00E-130 | 92 | 338 |
| YDRWdelta24 | 453 | 453 | 94% | 8.00E-130 | 92 | 338 |
| YDRWdelta23 | 453 | 453 | 94% | 8.00E-130 | 92 | 338 |
| YLRCdelta3 | 448 | 448 | 94% | 4.00E-128 | 92 | 338 |
| YERCdelta24 | 448 | 448 | 94% | 4.00E-128 | 92 | 338 |
| YLRCdelta18 | 448 | 448 | 97% | 4.00E-128 | 91 | 333 |

**Table S6.** Sequence similarity and identity search for *YDRWdelta23*.

| **Name** | **Max Score** | **Total Score** | **Cover** | **E value** | **Per. ident** | **Acc. Len** |
| --- | --- | --- | --- | --- | --- | --- |
| YPRWdelta21 | 625 | 625 | 100% | 0 | 100 | 338 |
| YPRWdelta20 | 625 | 625 | 100% | 0 | 100 | 338 |
| YMRCdelta9 | 625 | 625 | 100% | 0 | 100 | 338 |
| YMRCdelta10 | 625 | 625 | 100% | 0 | 100 | 338 |
| YLRCdelta2 | 625 | 625 | 100% | 0 | 100 | 338 |
| YLRCdelta3 | 619 | 619 | 100% | 7.00E-180 | 100 | 338 |
| YERCdelta24 | 619 | 619 | 100% | 7.00E-180 | 100 | 338 |
| YDRWdelta24 | 619 | 619 | 100% | 7.00E-180 | 100 | 338 |
| YOLWdelta4 | 614 | 614 | 100% | 3.00E-178 | 99 | 338 |
| YPRCdelta24 | 597 | 597 | 95% | 3.00E-173 | 100 | 338 |
| YPRCdelta23 | 588 | 588 | 94% | 2.00E-170 | 100 | 338 |
| YOLWdelta5 | 588 | 588 | 94% | 2.00E-170 | 100 | 338 |
| YJRWdelta13 | 586 | 586 | 100% | 7.00E-170 | 98 | 338 |
| YJRWdelta11 | 586 | 586 | 100% | 7.00E-170 | 98 | 338 |
| YGRWdelta14 | 586 | 586 | 100% | 7.00E-170 | 98 | 338 |
| YGRWdelta13 | 586 | 586 | 100% | 7.00E-170 | 98 | 338 |
| YMLWdelta4 | 569 | 569 | 100% | 7.00E-165 | 97 | 332 |
| YHRCdelta15 | 569 | 569 | 100% | 7.00E-165 | 97 | 332 |
| YGRCdelta30 | 569 | 569 | 100% | 7.00E-165 | 97 | 332 |
| YGRCdelta29 | 569 | 569 | 100% | 7.00E-165 | 97 | 332 |
| YORWdelta14 | 568 | 568 | 99% | 3.00E-164 | 97 | 332 |
| YORWdelta13 | 568 | 568 | 99% | 3.00E-164 | 97 | 332 |
| YMLWdelta3 | 564 | 564 | 100% | 3.00E-163 | 97 | 332 |
| YHRCdelta16 | 564 | 564 | 100% | 3.00E-163 | 97 | 332 |
| YBRWdelta13 | 564 | 564 | 100% | 3.00E-163 | 97 | 332 |
| YBRWdelta12 | 564 | 564 | 100% | 3.00E-163 | 97 | 332 |
| YNLCdelta2 | 532 | 532 | 94% | 1.00E-153 | 97 | 332 |
| YNLCdelta1 | 532 | 532 | 94% | 1.00E-153 | 97 | 332 |
| YGRWdelta32 | 532 | 532 | 94% | 1.00E-153 | 97 | 332 |
| YMRCdelta8 | 527 | 527 | 94% | 5.00E-152 | 97 | 332 |
| YMRCdelta7 | 527 | 527 | 94% | 5.00E-152 | 97 | 332 |
| YKRCdelta12 | 527 | 527 | 94% | 5.00E-152 | 97 | 332 |
| YGRWdelta11 | 527 | 527 | 94% | 5.00E-152 | 97 | 332 |
| YBLWdelta10 | 527 | 527 | 94% | 5.00E-152 | 97 | 332 |
| YPRWdelta12 | 521 | 521 | 94% | 2.00E-150 | 97 | 332 |
| YGLWdelta7 | 521 | 521 | 94% | 2.00E-150 | 97 | 332 |
| YBLWdelta9 | 518 | 518 | 94% | 3.00E-149 | 96 | 334 |
| YHRCdelta3 | 516 | 516 | 94% | 1.00E-148 | 96 | 332 |
| YKRCdelta8 | 499 | 499 | 94% | 1.00E-143 | 95 | 332 |
| YCRWdelta10 | 497 | 497 | 94% | 4.00E-143 | 95 | 331 |
| YFRCdelta8 | 494 | 494 | 94% | 5.00E-142 | 95 | 332 |
| YCRCdelta6 | 494 | 494 | 94% | 5.00E-142 | 95 | 332 |
| YDRWdelta11 | 486 | 486 | 94% | 8.00E-140 | 95 | 331 |
| YORCdelta25 | 486 | 486 | 93% | 8.00E-140 | 95 | 332 |
| YDRWdelta26 | 483 | 483 | 92% | 1.00E-138 | 95 | 332 |

**Table S7.** Sequence similarity and identity search for *TyA Gag* gene.

| **Name** | **Max Score** | **Total Score** | **Cover** | **E value** | **Per. ident** | **Acc. Len** |
| --- | --- | --- | --- | --- | --- | --- |
| YHRCTy1-1 | 983 | 1198 | 100% | 0 | 99 | 6028 |
| YPLWTy1-1 | 1016 | 1016 | 100% | 0 | 100 | 5924 |
| YOLWTy1-1 | 1016 | 1016 | 100% | 0 | 100 | 5926 |
| YMLWTy1-2 | 1016 | 1016 | 100% | 0 | 100 | 5903 |
| YLRWTy1-3 | 1016 | 1016 | 100% | 0 | 100 | 5920 |
| YLRWTy1-2 | 1016 | 1016 | 100% | 0 | 100 | 5918 |
| YLRCTy1-1 | 1016 | 1016 | 100% | 0 | 100 | 5922 |
| YJRWTy1-1 | 1016 | 1016 | 100% | 0 | 100 | 5922 |
| YERCTy1-1 | 1016 | 1016 | 100% | 0 | 100 | 5924 |
| YDRWTy1-5 | 1016 | 1016 | 100% | 0 | 100 | 5918 |
| YDRCTy1-3 | 1016 | 1016 | 100% | 0 | 100 | 5493 |
| YPRCTy1-4 | 1005 | 1005 | 100% | 0 | 100 | 5926 |
| YPRWTy1-3 | 989 | 989 | 100% | 0 | 99 | 5929 |
| YORWTy1-2 | 989 | 989 | 100% | 0 | 99 | 5914 |
| YGRCTy1-3 | 989 | 989 | 100% | 0 | 99 | 5914 |
| YGRCTy1-2 | 989 | 989 | 100% | 0 | 99 | 5918 |
| YDRWTy1-4 | 989 | 989 | 100% | 0 | 99 | 5926 |
| YDRCTy1-2 | 989 | 989 | 100% | 0 | 99 | 5918 |
| YDRCTy1-1 | 989 | 989 | 100% | 0 | 99 | 5918 |
| YBRWTy1-2 | 989 | 989 | 100% | 0 | 99 | 5917 |
| YMRCTy1-4 | 983 | 983 | 100% | 0 | 99 | 5926 |
| YMLWTy1-1 | 983 | 983 | 100% | 0 | 99 | 5914 |
| YERCTy1-2 | 983 | 983 | 100% | 0 | 99 | 5727 |
| YPRCTy1-2 | 977 | 977 | 100% | 0 | 99 | 5918 |
| YJRWTy1-2 | 977 | 977 | 100% | 0 | 99 | 5922 |
| YGRWTy1-1 | 977 | 977 | 100% | 0 | 99 | 5926 |
| YARCTy1-1 | 977 | 977 | 100% | 0 | 99 | 5925 |
| YNLWTy1-2 | 928 | 928 | 100% | 0 | 97 | 5900 |
| YNLCTy1-1 | 889 | 889 | 100% | 0 | 96 | 5914 |
| YBLWTy1-1 | 601 | 601 | 100% | ######## | 86 | 5916 |
| YMRCTy1-3 | 590 | 590 | 100% | ######## | 86 | 5914 |
| YORCTy2-1 | 226 | 226 | 50% | 2.00E-60 | 82 | 5961 |
| YNLCTy2-1 | 226 | 226 | 50% | 2.00E-60 | 82 | 5297 |
| YLRWTy2-1 | 226 | 226 | 50% | 2.00E-60 | 82 | 5959 |
| YDRWTy2-3 | 226 | 226 | 50% | 2.00E-60 | 82 | 5955 |
| YDRCTy2-1 | 226 | 226 | 50% | 2.00E-60 | 82 | 5956 |
| YGRCTy2-1 | 224 | 224 | 49% | 9.00E-60 | 81 | 5961 |
| YORWTy2-2 | 220 | 220 | 50% | 1.00E-58 | 81 | 5959 |
| YLRCTy2-2 | 220 | 220 | 50% | 1.00E-58 | 81 | 5443 |
| YGRWTy2-2 | 220 | 220 | 50% | 1.00E-58 | 81 | 5951 |
| YFLWTy2-1 | 220 | 220 | 50% | 1.00E-58 | 81 | 5959 |
| YDRWTy2-2 | 220 | 220 | 50% | 1.00E-58 | 81 | 5959 |
| YCLWTy2-1 | 220 | 220 | 50% | 1.00E-58 | 81 | 5959 |
| YBLWTy2-1 | 220 | 220 | 50% | 1.00E-58 | 81 | 5959 |

**Table S8.** Codon optimization of *AtCB5SD-PgFADX, PgFAD2, PgOLE1* and *RnELO2.*

| **Name** | **Length** | **CAI** | **%G+C** | **%G+C(1)** | **%G+C(2)** | **%G+C(3)** | **Nc** |
| --- | --- | --- | --- | --- | --- | --- | --- |
| fadx_before | 1575 | 0.665 | 51.9 | 52.8 | 42.1 | 61 | 55.4 |
| fadx_after | 1575 | 0.891 | 41.3 | 45 | 41.9 | 37 | 29 |
| fad2_before | 1164 | 0.591 | 58.6 | 54.9 | 42 | 78.9 | 46.4 |
| fad2_after | 1164 | 0.939 | 36.5 | 44.1 | 41.8 | 23.7 | 26.8 |
| ole1_before | 1524 | 0.702 | 51 | 57.5 | 43.3 | 52.4 | 52.4 |
| ole1_after | 1524 | 0.896 | 41.5 | 46.7 | 43.3 | 34.4 | 27.9 |
| elo2_before | 804 | 0.676 | 48.5 | 44 | 32.1 | 69.4 | 44.8 |
| elo2_after | 804 | 0.842 | 40.3 | 33.2 | 32.5 | 55.2 | 29.8 |
